# Aggregation-induced emission dots assisted non-invasive fluorescence hysterography in near-infrared IIb window

**DOI:** 10.1101/2021.02.14.431069

**Authors:** Xiaoming Yu, Yanyun Ying, Zhe Feng, Ji Qi, Junyan Zheng, Yuhuang Zhang, Juan Liu, Jun Qian, Ben Zhong Tang, Dan Zhang

## Abstract

Uterine diseases seriously threaten the physical and mental health of women. The main principle, when clinicians adopt examinations, is to achieve efficient diagnosis without negative effect on the physical function including fertility. Hysterography in near-infrared (NIR) IIb window (1500-1700 nm) presents perceptibly enhanced signal to background ratio (SBR) and higher penetration capability compared with those beyond 1000 nm and 1300 nm, but lays down high requirements for the biosafety of fluorophores at the same time. Assisted by the biologically excretable aggregation-induced emission (AIE) dots, non-invasive NIR-IIb fluorescence hysterography visualized typical Y-shaped uteruses, real-time uterine peristalsis or the uterine lesions (mimetic disease statuses in clinic) in mouse models. Significantly, after intrauterine perfusion, the reproductive capacity was unimpaired via fertility assessment and histological analysis. This work could inspire some new ideas for non-invasive clinical diagnosis of uterine diseases and effectively promote the clinical translation of AIE dots.

## 1. Introduction

Uterus is a vitally important organ for a women thoughout her life. It is dynamic in the physiology of menstrual cycle, embryo implantation, pregnancy immune maintain, child birth, and important paracrine or local endocrine function before or after menopause. The negative effect of uterine diseases on the human physical and mental health is no longer a matter for debate. Among these diseases, uterine cavity lesions (submucous myoma, endometrial polyp or neoplasm), intrauterine adhesion and uterine perforation/rupture affect female psychological and physical health, bring about infertility, and even impose huge personal as well as societal healthcare burdens.^[1–3]^

Examinations of gynecologic and obstetric diseases are chosen prudently for clinicians especially when female patients have fertility requirements or are during pregnancy period. In clinical practice, there are three kinds of widely used imaging technology for the diagnosis uterine diseases: ultrasound (US) imaging, magnetic resonance imaging (MRI) and hysterosalpingography (HSG), yet they have their own deficiencies. US imaging, limited by its poor spatial resolution, is not ideal in detecting lesions with small acoustic impedance difference. Moreover, the accuracy of the results largely depends on the operating skill and experience of the examiner.^[4]^ MRI, another imaging method, is known for accurateness with superior soft-tissue contrast, large field of view and multiplanar capabilities.^[5]^ It is commonly used to assess adenomyosis, uterine fibroid and neoplasm, rather than intrauterine adhesion and tubal patency while these abnormalities occupy quite a large proportion of female infertility cases. The third one, HSG, can sketch the contours of uterine cavity as well as the fallopian tubes through the flow of contrast agent in the cavity. But, it is a radiographic evaluation with the risk of ionizing radiation and fat embolism during the intravasation of the oil contrast while it mainly focuses on the patency of fallopian tubes.^[6]^ Briefly, there is a lack of biosafe, high-resolution, low-risk, real-time and affordable medical imaging method for detecting uterine diseases. It should be noted that non-invasive optical imaging modality has been indispensable in biological and medical research.^[7–10]^ Recently, it has been proved that photon scattering and autofluorescence of biological tissues are extremely low in the spectral region from 1500 nm to 1700 nm, which is usually named as the near-infrared IIb (NIR-IIb) window.^[11]^ Thus, NIR-IIb fluorescence imaging obtains the advantages of high spatial resolution, deep penetration and desirable signal-to-background ratio (SBR) with bright prospects in biomedicine, including the unexplored applications in reproductive system imaging.^[12–17]^

The precise treatment of uterine diseases through image-guided technology with no side effects of fertility or uterine physiology as possible is what reproductive experts and gynecologists expect to achieve. However, the advanced NIR-IIb fluorescence imaging technique generally requires bright exogenous optical probes,^[18–25]^ which inevitably introduces the biosafety concerns. So far, even though several types of NIR‐IIb fluorescent probes have been developed, unfortunately, some intrinsic drawbacks of specific fluorophores hampered their application in bioscience. For instance, heavy metals in quantum dots (QDs) may cause systemic changes including bone, hematopoietic system, nervous system, and especially the reproductive system.^[26, 27]^ When treating female patients with unsafe probe labeling, the adverse effects including subfertility, infertility, early pregnancy loss and even fetal malformation might be caused.^[28, 29]^ With great biocompatibility, various kinds of aggregation-induced emmition (AIE) dots have been successfully utilized in deep-penetration bioimaging.^[30–44]^ Very recently, the design scheme of excretable fluorophore AIEgens has been proposed in our previous work.^[45]^ The long aliphatic chains were verified that could be conductive to the excretion of OTPA-BBT molecules encapsulated in the injected dots from animal body and equip the PEGylated dots with excellent biosafety. Besides, the synthesized organic dots were proved secure for mice and even non-human primates. With aggregation-induced emission (AIE) property, the PEGylated OTPA-BBT aggregates in aqueous dispersion could emit bright fluorescence even beyond 1500 nm, which possesses strong potential for further clinical applications in NIR-IIb window.

Herein, we put forward a novel non-invasive NIR-IIb optical diagnostic method for uterine diseases in female reproductive system. In this work, excretable OTPA-BBT dots were utilized for both hysterography and uterine angiography. The emission properties of the dots in the second near-infrared (NIR-II, 900-1700 nm) window were characterized and the good photostability in intrauterine lavage fluid was confirmed. Then, the performance comparison of fluorescence hysterography in the first near-infrared (NIR-I, 760-900 nm), NIR-II, near-infrared IIa (NIR-IIa, 1300-1400 nm) and NIR-IIb windows was carried out in the same mouse, and high-quality NIR-IIb fluorescence uterine imaging *in vivo* was highlighted. The change of the fluorescent fluid level in uterine cavity was observed along with the uterine peristalsis during the fluorescence hysterography. Subsequently, the uterine perforation/rupture and followed effective surgical repair in the mouse model were identified, simulating the clinical scene when it occurs in surgery. Next, the partial and complete uterine obstruction models in mice were established to demonstrate *in vivo* diagnosis of uterine malformation and intrauterine diseases via fluorescence hysterography. Eventually, the influence on fertility was evaluated after treatment. All the excellent performance presented the feasibility and effectiveness of AIEgens-based hysterography in NIR-IIb window as well as great application potential in further clinical practice.

## 2. Main text

### 2.1. Fabrication and characterization of OTPA-BBT dots

**Figure 1A** illustrates the chemical structure of OTPA-BBT molecule, whose AIE characteristic has been verified in our previous work. The molecular synthesis of OTPA-BBT is described in Scheme S1 and Figure S1-7. To hydrate the AIEgens, OTPA-BBT molecules were encapsulated into organic nanoparticles using US Food and Drug Administration (FDA)-approved F-127 (Figure 1B). As shown in Figure 1C and 1D, the absorption and photoluminescence (P.L.) spectra of OTPA-BBT dots in deionized (DI) water were measured. To minimize the autofluorescence, the 793 nm laser, whose wavelength is closed to but longer than the peak absorption wavelength, was selected as the excitation source. The fluorescence emission tail could extend even beyond 1500 nm (Figure 1D) and the quantum yield was as high as 13.6% in NIR-II region.^[45]^ As expected, the organic dots presented the considerable brightness in NIR-IIb region (Figure 1E). The image taken via scanning transmission electron microscope (STEM) in Figure S8A illustrated the synthesized OTPA-BBT dots possessed uniform sphere structure. Besides, these dots had an average diameter of 28.3 ± 1.6 nm from dynamic light scattering (DLS), as shown in Figure 1F and Figure S8B.

**Figure 1.**
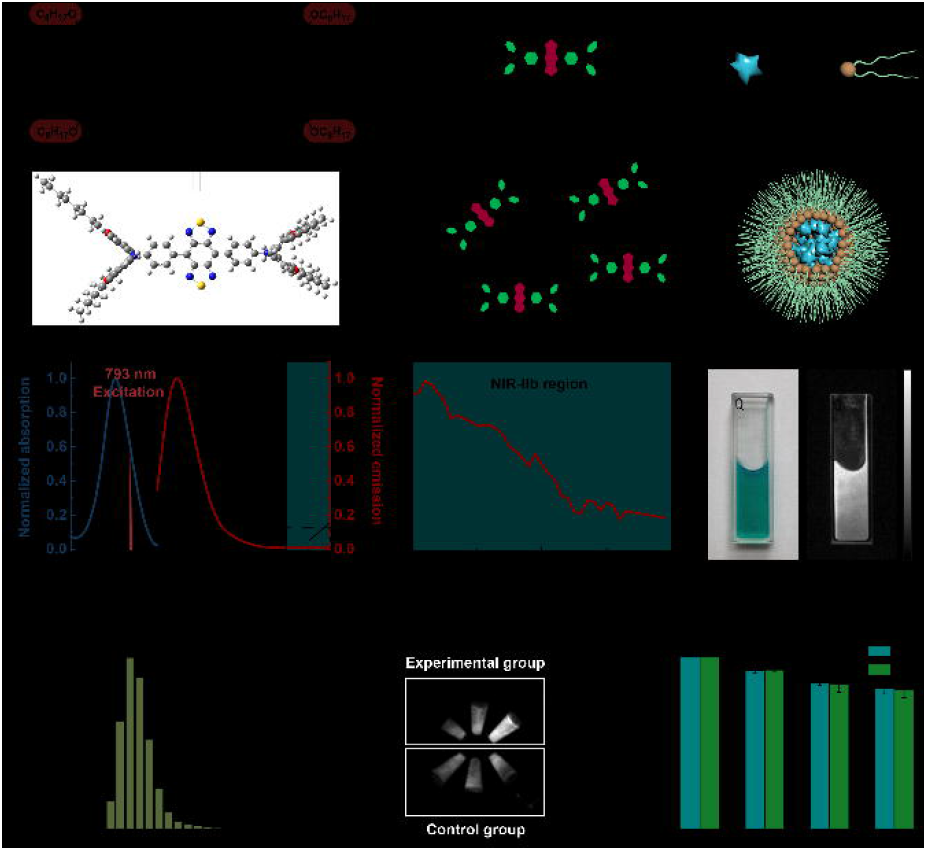
The synthesis and characterization of OTPA-BBT dots. A) Molecular structure of OTPA-BBT. B) Schematic illustration of the fabrication of OTPA-BBT dots. C) Normalized absorption and emission spectra of OTPA-BBT dots. D) The local enlarged PL spectrum of OTPA-BBT dots beyond 1500 nm. E) Bright-field picture and NIR-IIb fluorescence image of OTPA-BBT dots in water. F) Representative DLS result of the OTPA-BBT dots. G) NIR-IIb fluorescence images of OTPA-BBT dots dissolved in DI water and intrauterine lavage fluid (n = 3). The intra group difference of the brightness was caused by the concentration difference. H) Fluorescence stability of OTPA-BBT dots irradiated with 793 nm laser (power density, 120 mW cm^−2^) for 30 min.

Good photostability in the uterine cavity is the prerequisite for the hysterography, thus we collected intrauterine lavage fluid for *in vitro* test. As shown in Figure 1G and 1H, the NIR-IIb emission intensity of OTPA-BBT dots dilutions in intrauterine lavage fluid had changed little compared to that in DI water at the same condition, under continuous high-intensity illumination (~120 mW cm^−2^) of 793 nm continuous wave (CW) laser for 30 min. The results also proved that the emission of dots was unaffected much in the certain biotic environment. The morphologies of OTPA-BBT dots in strong acidic (pH = 2) and alkaline (pH = 12) solutions were observed as displayed in Figure S9 which further confirmed the excellent chemical stability. Considering the convenience of preparation and long-term storage as future clinical medicament, the OTPA-BBT dots were freeze-dried and then rehydreated. The freeze-dried OTPA-BBT dots possessed excellent water solubility (Figure S10A, B). The NIR-IIb fluorescence images in the Figure S10C, D showed that freeze-dried and rehydrated OTPA-BBT dots both had good brightness beyond 1500 nm, and the normalized absorption spectra of the OTPA-BBT dots before freeze-drying and after rehydration were also highly coincident (Figure S10E). Not only that, the STEM image and DLS result could be the verification for the stable morphology and particle size (Figure S10F, G). The verified high stablity, convenient storage and preparation, as well as the excellent biosafty demonstrated in our previous work,^[45]^ made the OTPA-BBT dots contain great potential for gynecological diagnosis.

### 2.2. Fluorescence imaging of the reproductive system

The priority of revealing the genital system is to choose minimally-invasive or non-invasive imaging methods, rather than the invasive ones. The current diagnostic technology for uterine diseases in clinic, such as HSG, might expose reproductive organs to ionizing radiation and make the patients face the risk of allergy to radiographic lipiodol contrast. Therefore, the exploration of novel diagnostic technology is essential and the NIR-IIb fluorescence imaging could be a promising candidate.

In order to closely match the simulated scenes in clinical practice, NIR-IIb fluorescence visualization of the uterus *in vivo* in mouse model was applied. As shown in **Figure 2A**, with the assistance of the bright and biosafe OTPA-BBT dots, hysterography (the morphology of the uterine cavity) and angiography (the outline of the uterine vessels) could be conducted via intrauterine perfusion and intravenous injection, respectively.

**Figure 2.**
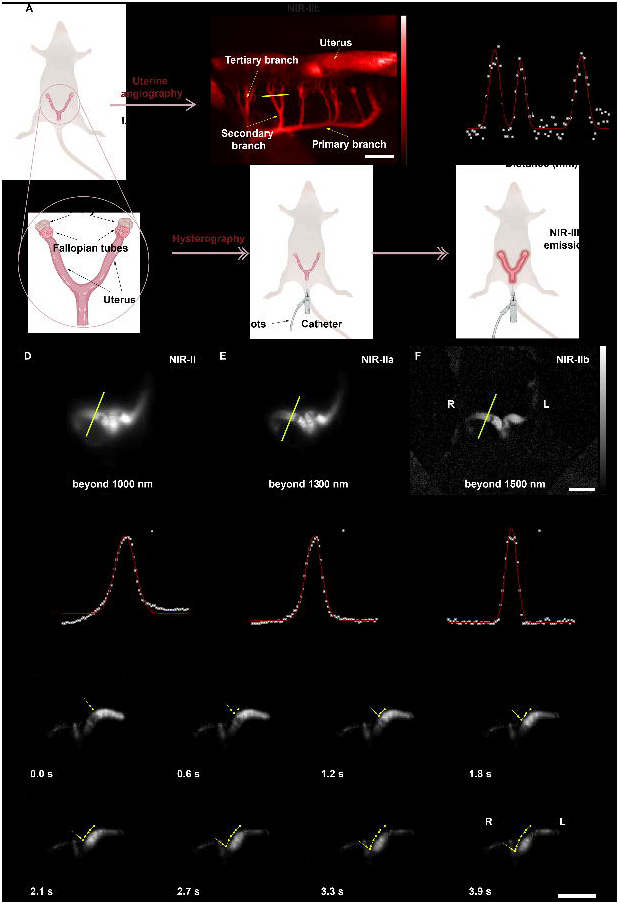
Both hysterography and angiography in NIR-IIb window of female mice. A) The illustration of the anatomy structure and the position of female genital system, the modes of administration as well as the expected imaging results. B) Uterine vessels in left side was imaged in the NIR-IIb window. Scale bar, 1.25 mm. C) A cross-sectional fluorescence intensity profile along the yellow solid bar of tertiary branches (inset of Figure 2B). The Gaussian fit to the profile was shown as the red line. Representative fluorescence images of the Y-shaped uterus after intrauterine perfusion of OTPA-BBT dots beyond D) 1000 nm, E) 1300 nm and F) 1500 nm. The power densities of the laser irradiation were ~10, ~80, ~80 mW cm^−2^, respectively. The integrated times were 5, 15, 150 ms, respectively. Scale bar, 10 mm. G-I) The FWHMs and SBRs of the uterine imaging in various optical tissue windows were calculated by Gaussian fits to the intensity profiles along the yellow solid bars in Figure D-F. J) *In vivo* NIR-IIb fluorescence imaging of uterine peristalsis was reflected by the change of the fluid level (yellow line) in the cavity at various time points perfused with OTPA-BBT dots. Yellow arrows indicated the current fluid levels and yellow dots indicated the fluid levels in the past. Scale bars, 20 mm. L, left; R, right.

#### 2.2.1. The uterine angiography in NIR-IIb window

Recently, NIR-IIb spectral region has been verified as an excellent optical tissue window for vessel imaging *in vivo*. Due to the reduced light scattering^[46, 47]^ and restrained autofluorescence^[12]^, the tertiary branches of uterine vessels in the unilateral uterus were vividly presented under the excitation of 793 nm CW laser (~80 mW cm^−2^) after intravenous injection of OTPA-BBT dots (1 mg mL^−1^, 200 μL) (Figure 2B). The analysis of uterine angiography quality was presented in Figure 2C. The full widths at half-maximum (FWHMs) of the fitting curves were calculated as 220 μm, 180 μm and 240 μm, respectively, exhibiting high spatial resolution. As shown in Figure S11, the excretion of OTPA-BBT dots could be confirmed, which is consistent with the conclusion in our previous work and the excretion can be attributed to the long aliphatic chains.

#### 2.2.2. Uterine imaging in NIR-II/NIR-IIa/NIR-IIb windows in mice without laparotomy

Visualization of the uterus always needs to penetrate the skin, subcutaneous fatty layer and muscle layer as well as perimetric fat pad. Precise deciphering of bio-tissue at large depth equips NIR-IIb fluorescence imaging with great potential for non-invasive hysterography *in vivo*. In our work, typical Y-shaped uterus of the mouse was visualized via uterine perfusion of OTPA-BBT dots beyond 1000 nm (Figure 2D). By converting the long-pass filters, the fluorescence images of the uterus beyond 1300 nm and 1500 nm were also taken (Figure 2E, F). As the increase of the detection wavelengths, the images exhibited enhanced contrast and less stray light signals, significantly sharping the contour of uterine cavity. By quantitative calculation, the FWHMs of the fitting curves in Figure 2D, 2E and 2F were calculated as 2.02 mm, 1.66 mm and 1.16 mm, respectively. SBR was calculated as 115.03 in the NIR-IIb window, which was much higher than those measured in the NIR-II/NIR-IIa windows (8.99 and 23.63, respectively) (Figure 2G-I). In addition, to futher simulate intraoperative and postoperative diagnosis, we performed uterine fluorescence imaging in various optical windows before and after the mouse abdomen closed. As presented in Figure S12, there was no doubt that fluorescence imaging in NIR-IIb region was superior to the others.

On the other hand, uterine peristalsis, originating from the contraction of the myometrium, performs as the uterine cavity pressure changing and endometrial movement, whose frequency and direction influence numerous steps of life activities (such as menstruation, fertilization, embryo implantation and fetal delivery).^[48–50]^ In addition to sharp spatial resolution, NIR-IIb fluorescence imaging could also show the dynamic flow of fluorescent solution in the uterine cavity with the strength of high temporal resolution. The fluid level declined along with uterine peristalsis from the top to the bottom position. As shown in Figure 2J and Movie S1, the record of the certain position of the fluid level at each observing time point well presented its track, reflecting the real-time physical state of the uterus. As a tubal structure, fallopian tubes also have the physical characteristic of directional peristalsis. This imaging method might give a novel approach to assist the diagnosis of tubal factor infertility in the further clinical applications.

### 2.3. NIR-IIb fluorescence hysterography for uterine perforation/rupture identification and repair

In clinic, intrauterine operations, including myomectomy, adenomyosis focus resection, uterine septum resection and uterine deformity reduction are always accompanied by the risk of uterine perforation and rupture, which is traditionally not easy to be diagnosed precisely and instantly. However, if the perforation is not repaired in time, it might result in further bleeding, infection, and even bowel prolapse and perforation. But at present, there is neither convenient nor effective approach to detect uterine perforation/rupture and to facilitate prompt clinical treatment strategy.

Given the superiority of imaging window from 1500 nm to 1700 nm, the NIR-IIb fluorescence hysterography was applied to the identification of uterine perforation/rupture and followed repair in a mouse model (**Figure 3A**). As displayed in Figure 3B and 3C, the middle position of the left uterus was incised with surgical scissors and then the abdominal incision was sutured. As shown in Figure 3D and Movie S2, after the uterine perfusion of OTPA-BBT dots, the whole process of the solution leaking from the left uterus was recorded in time through real-time NIR-IIb fluorescence imaging. Initially, the outline of the uterus began to appear from the bottom position to the middle position. Notably, at 3.4 s the fluorescent solution leaked into the abdominal cavity through the surgical incision and the size of leakage area increased rapidly with time. The leakage area raised from ~100 pixels to ~820 pixels as the time went from 3.4 to 4.6 s (Figure 3E). Thus, the excellent imaging performance in NIR-IIb window showed superb potentialities for clinical translation to accurately determine the occurance of the uterine perforation/rupture. Subsequently, the defect was ligated using a 5-0 suture and the abdominal cavity was flushed with 37 °C 1× phosphate-buffered saline (PBS) after the abdominal cavity was opened (Figure 3F, G). After half an hour, no obvious fluorescence signal was seen as the fluorescent dots flowed away along the cervix and vagina (Figure S13). After perfusion again, fluorescence hysterography was performed under the excitation of 793 nm CW laser. As shown in Figure 3H, 3I and Movie S3, there was no leakage of fluorescent solution from the original defect site in a period of time, which illustrated the effective surgical repair.

**Figure 3.**
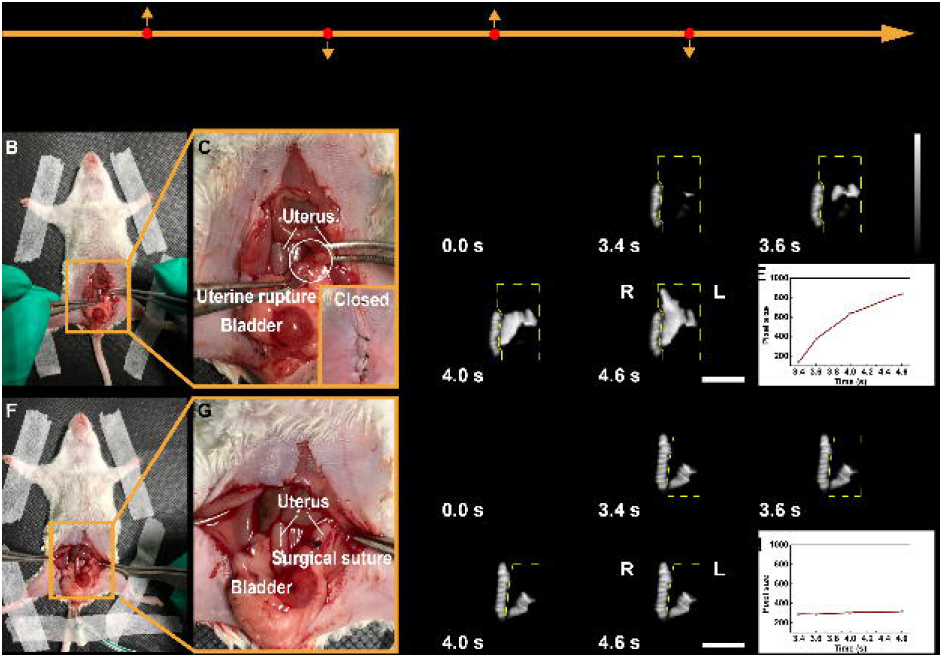
Identification of uterine perforation/rupture and followed effective suture in a mouse model via NIR-IIb fluorescence hysterography. A) The road map to achieve abovementioned procedure. B, C) The physical map of uterine rupture in a mouse model. D) Dynamic hysterography after intrauterine perfusion. E) Before the repair of uterine rupture, the size of leakage of OTPA-BBT dots. F, G) The physical map of the suture of uterine rupture in the mouse model. H) Dynamic hysterography after intrauterine perfusion. I) After the repair of uterine rupture, the size of leakage of OTPA-BBT dots. Scale bar, 10 mm. L, left; R, right. The power density of the laser irradiation was ~80 mW cm^−2^ and the integrated time was 150 ms.

### 2.4. NIR-IIb fluorescence hysterography for diagnosis of complete/partial uterine obstruction

Some intrautrine lesions, including uterine septum, adhesions, polyps, etc., directly affect the intrauterine environment and threaten the health of female patients and the survival of fetuses. Therefore, the precise positioning via hysterography could help us diagnose the uterine obstruction or stenosis and guide subsequent treatment.

In this study, complete/partial uterine obstruction models were established in mice, which mimicked the alteration of cavity induced by the above diseases. **Figure 4A** illustrated the pretreatment that the right uterus was ligated by utilizing a 5-0 suture while the left one received no treatment. After that, the abdominal incision was sutured. Consisted with our expectation, the non-treated side was imaged with full length after the uterine perfusion (Figure 4B). Meanwhile, the liquid flow was blocked in the lesion side and no fluorescence signal was observed above the ligation point. After the relief of obstruction and the close of abdomen, the uterus was imaged integrally no matter in the non-treated side or the lesion side under the same imaging condition (Figure 4C, D). Subsequently, another mouse model of partial obstruction in right uterus was built by placing its own adipose tissue into the cavity (Figure 4E). After perfusion of the OTPA-BBT dots, the right uterus above the side of fat tamponade was delayed to light up and off compared with the left uterus (Figure 4F), which we called the “delayed effect”, and the larger flowing resistance was indicatd in the lesion side due to foreign body. The OTPA-BBT dots then flowed away from the body via natural orifice and no obvious vestigial fluorescence signal could be detected. Before the abdomen was closed, the foreign body was taken out (Figure 4G). The “delayed effect” was not observed after uterine perfusion once again and the flowing resistance in bilateral uterine cavities varied slightly when the solution was injected and pumping back (Figure 4H). In the evaluation of tubal patency by chromopertubation (instillation of methylene blue through the fallopian tubes), peritoneal lavage is not an indispensable step because methylene blue has good biological safety. Thus, even if the OTPA-BBT dots in the abdominal cavity could not be removed completely by peritoneal lavage, introducing some extra background, they did not cause obvious damage thanks to their good biosafety. In clinical practice, the incidence of congenital malformation and dysplasia of the uterus could happen. Thus, we further established unilateral uterine defect model. As shown in Figure S14, signals were detected in the left area of the abdomen but missing in the opposite. Precise positioning of lesion site was satisfactorily presented in NIR-IIb window, which was greatly helpful to the diagnosis of uterine anomalies and intrauterine lesions.

**Figure 4.**
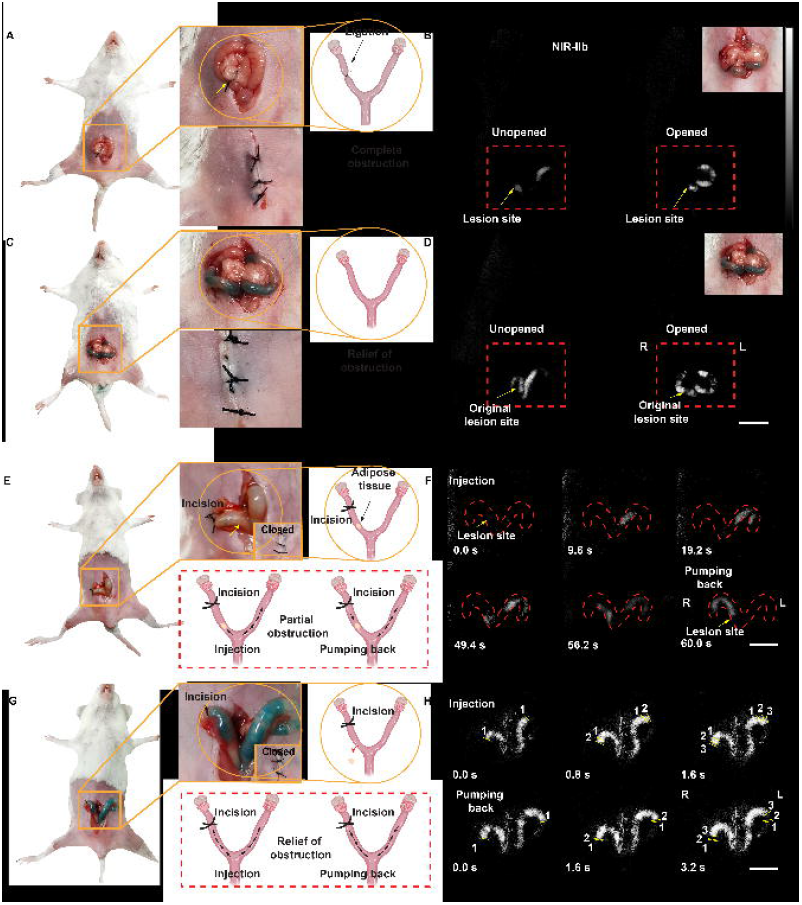
Detection of the partial and complete uterine obstruction in mouse models via NIR-IIb fluorescence hysterography. A) The physical map and the illustration of the complete uterine obstruction by ligation in a mouse model. B) NIR-IIb fluorescence hysterography through intrauterine perfusion before and after opening the abdomen. C) The physical map and the illustration of the relief of uterine obstruction by releasing the ligation. D) NIR-IIb fluorescence hysterography through intrauterine perfusion before and after opening the abdomen. E) The physical map of the partial uterine obstruction by intrauterine autologous fat filling in a mouse model. The “delayed effect” illustrated the unequal pressure of the uterine cavity caused by the foreign mass during perfusion. F) The “delayed effect”: NIR-IIb fluorescence hysterography via intrauterine perfusion after closing the abdomen. The signal delayed to appear and disappear in the right side G) The physical map of uterine obstruction by taking out the autologous fat and the illustration of the disapperance of the “delayed effect”. H) NIR-IIb fluorescence hysterography via injection and pumping back. Yellow lines showed liquid level at different time as pressure changed in uterine cavity. Scale bar, 10 mm. L, left; R, right. The power density of the laser irradiation was ~80 mW cm^−2^ and the integrated time was 150 ms.

### 2.5. Fertility assessment and histological analysis after intrauterine perfusion of OTPA-BBT dots

Due to scientists’ ceaseless efforts, many optical probes with strong NIR-IIb emission have continually emerged. However, the consequent concerns about biosafety of exogenous fluorophores have been increasing with time. Infertility affects arround 10% of reproductive-age women, and many factors contribute to this heterogeneous disease, including heavy metals, environmental pollutants, etc. It is a regret that some NIR-IIb fluorescent materials use reproductive toxic components such as heavy metals in order to guarantee sufficient brightness. Recently, our work has illustrated the good biosafety of OTPA-BBT dots via intravenous injection in rodent and non-human primate models by pathologic study, biochemical and metabolism analysis. Whereas, it is much more significant for women of childbearing age to remain their fertility unaffected, and the fertility assessment should be list as another necessary issue on the biocompatibility of OTPA-BBT dots.

Therefore, 16-week-old male ICR mice with proven fertility were utilized to mate with female mice to evaluate the influence of fertility after intrauterine perfusion of OTPA-BBT dots. There was no significant difference in duration from mating to first birth and the first litter size between the control group (n = 9, mean litter size = 12.44 ± 0.97, mean days = 26.56 ± 0.85) and experiment group (n = 9, mean litter size = 12.44 ±1.30, mean days = 25.56 ± 1.07), as displayed in **Figure 5A-D**. The results showed that the treatment with OTPA-BBT dots had no significant effect on the fertility of female mice. In order to observe the residue of OTPA-BBT dots in the reproductive system and the distribution, we performed NIR-II fluorescence imaging on ovaries, uteruses and fallopian tubes of mice at 1 day and 30 days post intrauterine perfusion. No fluorescence signal was observed in the female reproductive organs which were dissected out and imaged beyond 1300 nm at 1 day and 1 month post treatment (Figure 5E). Besides, no NIR-II fluorescent dots (beyond 1300 nm) remained in the main organs, which was demonstrated by bio-distribution test (Figure S15). These results proved that the OTPA-BBT dots could hardly be absorbed and accumulated in the reproductive system. The histopathologic observation and analysis were then performed. As shown in Figure 5F, compared with the control group, the endometrial thickness, tubal villi and ovarian structure of the experimental group did not change significantly. Thus, biosafe and fertility-friendly OTPA-BBT dots are desirable for NIR-IIb fluorescence hysterography, which could be applied to the examination for women in childbearing age.

**Figure 5.**
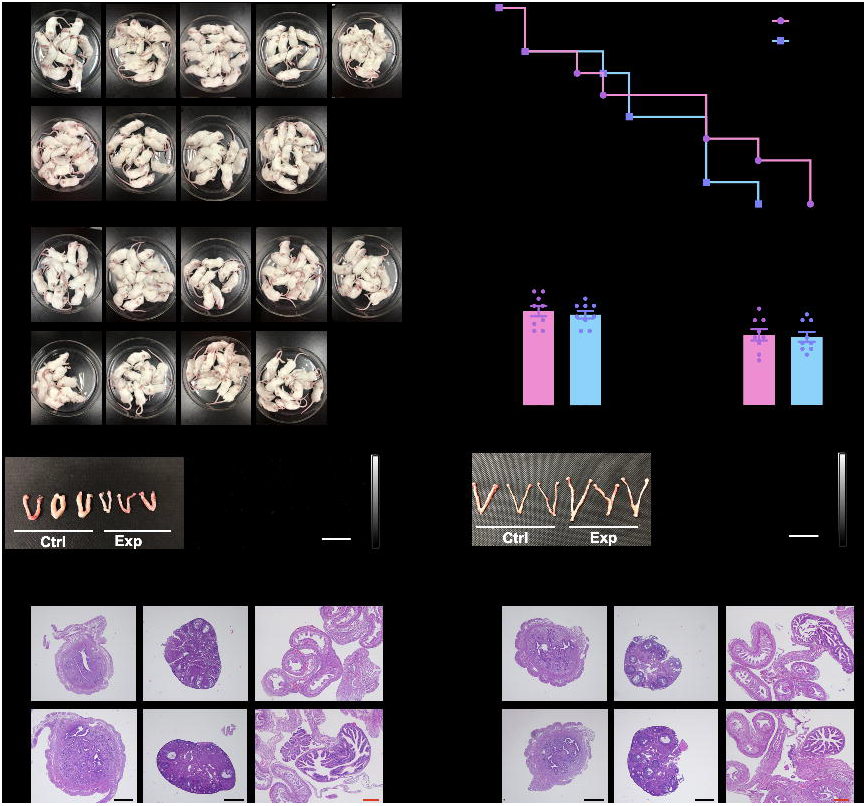
The fertility assessment and reproductive toxicity analysis after intrauterine perfusion of OTPA-BBT dots. A) The offspring amount of each mouse in control group and experiment group (n = 9). B) Mating days until first birth. X, The time from mating to labor. Y, Number of pregnant mice. C) The mean duration from mating to first birth. D) The mean litter size of first litter after treatment. E) The NIR-II fluorescence images of genital system at 1 day and 1 month after treatment. Scale bar, 30 mm. F) Representative H&E stain of uteruses, ovaries and fallopian tubes. Black scale bar, 500 μm; Red scale bar, 200 μm. Error bars represent SEM.

## 3. Conclusion

Generally, the diagnosis and treatment applied in reproductive system imaging are required to be non-invasive and non-toxic, with minimized side effect on fertility and other physical function, which set high standards on the biosafety of theranostics. Nowadays, non-invasive NIR-IIb fluorescence imaging with high contrast and spatial resolution has been verified to possess great potential for clinical translation. In addition, the OTPA-BBT dots were proved to be excretable when injected intravenously into animals in our previous work, indicating that the biosafe dots could greatly reduce the risk of toxicity in case the fluorophores entered blood during the operation. Moreover, OTPA-BBT dots exhibited bright NIR-IIb emission. Thus, we believed the OTPA-BBT dots assisted NIR-IIb fluorescence imaging could be a novel approach for hysterography in female patients.

In this work, OTPA-BBT dots were utilized for both angiography and hysterography in NIR-IIb window. Through fluorescence angiography, we observed the tertiary branches of uterine vessels with satisfying spatial resolution and contrast. NIR-IIb fluorescence hysterography showed preferable performance when outlining the uteruses as well as natural peristalsis. Moreover, real-time visualization ability enabled NIR-IIb fluorescence imaging to judge the occurance of the uterine perforation, which is one of the common complications in hysteroscopic surgery. The uterine obstruction and stenosis, caused by uterine lesions like severe intrauterine adhesion, were easily recognized via NIR-IIb fluorescence imaging. It was helpful to locate the lesion site for further precise treatment in above clinical scenes. Notably, there was almost no influence of the OTPA-BBT dots upon the fertility after the assessment in duration from mating to first birth and the number of first litter size. Non-invasive and fertility-friendly fluorescence hysterography in NIR-IIb region was achieved for the first time, which might be able to guide the trend of examinations especially for female patients in childbearing age.

So far in clinical practice, the severity of uterine perforation is still unable to be accurately measured by imaging methods. With high spatial and temporal resolution of fluorescence hysterography, we would be able to calculate volume leakage flow to evaluate the defect scale by computer modeling in the future, which might be achieved by multi-dimensional fluorescence imaging. In our work, the uterine peristalsis was clearly observed. While fallopian tubes possess the similar physiological characteristics of directional peristalsis. Theoratically, the fluorescence imaging could be extended to dynamically determine the tubal patency and pathological peristalsis. Nevertheless, fallopian tubes could not be presented owing to the special structure in the mouse, so that this aspect could not be verified in mouse model. Therefore, applying fluorescence tubal imaging into appropriate model animals has been incorporated into our plan.

## 4. Methods

### Molecular synthesis

The detailed synthesis process of OTPA-BBT is available in the Supporting Information.

### Materials

Pluronic F-127 was purchased from Sigma-Aldrich. DI water prepared by an Eco-Q15 deionized water system (Shanghai Hitech Instruments Co., Ltd.) was used in all the experimental procedures. PBS was obtained from Sinopharm Chemical Reagent Co., Ltd., China. The tetrahydrofuran (THF) was purchased from Shanghai Aladdin Biochemical Technology Co., Ltd., China.

### *PEG*ylation of AIEgens

2 mg OTPA-BBT AIEgens and 10 mg Pluronic F-127 were dissolved in 1 mL THF. Then, the total solution was added into 10 mL DI water. Next, the mixture was sonicated with a microtip probe sonicator (XL2000, Misonix Incorporated, NY) for nearly 5 min. The colloidal dispersion (OTPA-BBT dots) was obtained by stirring the suspension violently overnight in the fume hood, in which THF solvent was completely evaporated.

### Absorption and fluorescence emission spectra mesurement

The absorption spectrum of the OTPA-BBT dots in water was recorded by a Shimadzu UV-2550 UV–vis–NIR scanning spectrophotometer. The fluorescence emission spectrum of OTPA-BBT dots in water in the NIR-II window was measured by a lab-built system (Figure S16) based on a NIR2200 spectrometer (Ideaoptics Instruments).

### Animal preparation and ethical approval

Female Institute of Cancer Research (ICR) mice (6-8 weeks old) and male ICR mice (16 weeks, for mating) were obtained from the Shanghai SLAC Laboratory Animal Co.,Ltd (Shanghai, China). The animals were housed at 21-24 °C, relative humidity of 40-60% with a 12 h light/dark cycle and fed with standard rodent laboratory chow and water. All experimental protocols in this study were performed in accordance with the guidelines for the humane use of laboratory animals and were approved by the Ethics Committee of the Laboratory Animal Center of Zhejiang University (reference number, ZJU20200120). Anesthesia was used for all the animals prior to and during the experimental interventions.

### NIR-IIb fluorescence in vivo uterine angiography

Female ICR mice were anesthetized with 1.25% tribromoethanol (Sigma-Aldrich) (250 mg kg^−1^) by intraperitoneal injection. Then the mouse abdomen was shaved and depilated before experimental interventions. Under anesthesia, mice’ limbs were fixed on a flat with medical aerated tapes (3M), and then a lower midline vertical incision (around 2 cm) in the abdomen was made. After that, the mice were kept in lateral position. The left uterus was isolated, gently pulled out with forceps and fixed by stainless steel pins to another platform when uterine vessels were fully exposed. The region of interest was placed under a lab-built NIR-II fluorescence whole-body imaging system (Figure S17) aiming to visualize the uterine vessels and their collateral branches. Under the excitation of 793 nm CW laser with the power density of ~80 mW cm^−2^, the NIR-IIb fluorescence signals were collected by a fixed focus lens (with near infrared antireflection) through a 1500 nm long-pass optical filter, after intravenous injection of OTPA-BBT dots (1 mg mL^−1^, 200 μL).

### Non-invasive NIR-IIb fluorescence hysterography without a laparotomy

Female ICR mice were anesthetized with 1.25% tribromoethanol (250 mg kg^−1^) by intraperitoneal injection. The abdomen was shaved and depilated before experimental interventions. The female mice (non-treated or uterine disease models) were then placed in a supine position under the InGaAs camera to visualize the uterus. 200 μL OTPA-BBT dots were infused into the uterine cavity through cervix using a 26 G I.V. catheter (Jiangxi Fenglin, China). Under the excitation of 793 nm CW laser with power density of ~80 mW cm^−2^, NIR-IIb fluorescence of OTPA-BBT dots perfused into the uterus was detected by the macroscopic imaging system.

### The establishment of uterine perforation/rupture and repair model in mice

Mice were anesthetized for surgery via 1.25% tribromoethanol (250 mg kg^−1^) by intraperitoneal injection. After animal preparation, laparotomy was performed and a small incision was made near the junction of left uterus and oviduct. Next, abdominal incision was closed with 5-0 silk surgical sutures. NIR-IIb fluorescence imaging was performed to visualize the leakage of OTPA-BBT dots during intrauterine perfusion. The leakage area was described as the occupied pixels, which was lighted up around the left uterus. After imaging, abdominal cavity was opened again, and uterine incision was stitched up. Peritoneal lavage was performed by 37 °C 1× PBS to remove the fluorescent agent. The abdominal cavity was stitched up until no significant signal could be detected by the InGaAs camera. Eventually, NIR-IIb fluorescence hysterography was performed to determine whether the uterus was repaired successfully.

### The establishment of complete and partial uterine obstruction models in mice

The complete uterine obstruction model was established by uterine ligation. Under anesthesia, the mouse was secured to a platform in the supine position, and then laparotomy was performed. The right uterus was isolated, fully exposed and then occluded at the lower 1/3 site by using a 5-0 silk surgical suture. Abdominal incision was closed and next NIR-IIb fluorescence uterine imaging was performed to detect the site of stenosis or obstruction. On the basis of the complete obstruction model, removing the right uterus from the distal part of the ligation site simulated the absence of uterus.

The partial obstruction model was established by intrauterine autologous fat filling. First, a small incision was made in the right uterus near the junction of uterus and oviduct to fill in the adipose tissue at the lower 1/3. Then, the position lower of the incision was ligated to seal the uterine cavity. Next, the abdominal cavity was closed for hysterography. At last, the fluorescence hysterography was performed again after adipose tissue was removed through original incision.

### Fertility assessment

Eighteen female ICR mice (8 weeks old) were randomly assigned into the experiment group and the control group. Anesthetized with 1.25% tribromoethanol, each mouse in the experiment group was given intrauterine perfusion with 200 μL OTPA-BBT dots (1 mg mL^−1^) and each mouse in the control group was given intrauterine perfusion with 200 μL PBS solution (1×) by catheter. After treatment, female mice (n = 9 for each group) were continuously mated with 16-week-old fertile males (female: male = 2:1) over 1 month. The time of labor and the number of pups were recorded.

### Histological analysis of genital system

At 1 day and 1 month post intrauterine perfusion, female mice were sacrificed and the organs including heart, liver, spleen, lung, kidney, ovary, uterus and fallopian tubes were collected. The ovaries, uteruses and fallopian tubes were fixed in 10% formalin, dehydrated in graded alcohol and xylene, and embedded in paraffin. Paraffin-embedded samples were sectioned at 5 μm thickness and stained with hematoxylin and eosin (H&E) for morphological observation. Pathology sections were captured under an optical microscope (Olympus BX61).

### Statistical analysis

Statistical analysis was performed with GraphPad Prism software version.8.2.1 (GraphPad Software, San Diego, CA). Results were given as means and standard error of means (SEMs). Group comparisons were made by two-tailed unpaired Student’s t-tests. P-values of less than 0.05 were considered to be statistically significant.

## Supporting Information

Supporting Information is available from the Wiley Online Library or from the author.

## Acknowledgements

X.Y., Y.Y, Z.F., J.Q. contributed equally to this work. This work was supported by National Key Research and Development Program of China (2018YFC1005003), National Natural Science Foundation of China (61735016, 81771535 and 61975172), Key Research and Development Program of Zhejiang Province (2021C03098), Zhejiang Provincial Natural Science Foundation of China (LR17F050001, WKJ-ZJ-1826), and Fundamental Research Funds for the Central Universities (2020-KYY-511108-0007). The procedures for all the animal experiments were approved by Animal Use and Care Committee at Zhejiang University (ZJU20200120) and in accordance with the National Institutes of Health Guidelines.

We are grateful to Jianhua Chen from Department of Pathology, Women’s Hospital, Zhejiang University School of Medicine for his technical guidance on tissue sections and HE staining. We thank Dandan Song in the Center of Cryo-Electron Microscopy (CCEM), Zhejiang University for her technical assistance on Scanning Transmission Electron Microscopy.

Received: ( )

Revised: ( )

Published online: ( )

## References

[1] E. Taylor, V. Gomel, Fertil. Steril. 2008, 89, 1.

[2] C. Massarotti, G. Gentile, C. Ferreccio, P. Scaruffi, V. Remorgida, P. Anserini, Gynecol. Endocrinol. 2019, 35, 485.

[3] B. W. Rackow, E. Jorgensen, H. S. Taylor, Fertil. Steril. 2011, 95, 2690.

[4] F. Schwab, K. Redling, M. Siebert, A. Schotzau, C. A. Schoenenberger, R. Zanetti-Dallenbach, Ultrasound in Medicine and Biology 2016, 42, 2622.

[5] T. Fukunaga, S. Fujii, C. Inoue, N. Mukuda, A. Murakami, Y. Tanabe, T. Harada, T. Ogawa, Jpn. J. Radiol. 2017, 35, 697.

[6] O. Uzun, S. Findik, M. Danaci, D Katar, L. Erkan, Respirology 2004, 9, 134.

[7] G. D. Luker, K. E. Luker, J. Nucl. Med. 2008, 49, 1.

[8] G. Hong, J. C. Lee, J. T. Robinson, U. Raaz, L. Xie, N. F. Huang, J. P. Cooke, H. Dai, Nat. Med. 2012, 18, 1841.

[9] Q. Wang, Y. Dai, J. Xu, J. Cai, X. Niu, L. Zhang, R. Chen, Q. Shen, W. Huang, Q. Fan, Adv. Funct. Mater. 2019, 29, 1901480.

[10] G. Hong, J. T. Robinson, Y. Zhang, S. Diao, A. L. Antaris, Q. Wang, H. Dai, Angew. Chem. Int. Ed. 2012, 51, 1.

[11] S. Diao, J. L. Blackburn, G. Hong, A. L. Antaris, J. Chang, J. Z. Wu, B. Zhang, K. Cheng, C. J. Kuo, H. Dai, Angew. Chem. Int. Ed. Engl. 2015, 54, 14758.

[12] S. Diao, G. Hong, A. L. Antaris, J. L. Blackburn, K. Cheng, Z. Cheng, H. Dai, Nano Res. 2015, 8, 3027.

[13] Z. Ma, F. Wang, W. Wang, Y. Zhong, H. Dai, Proc. Natl. Acad. Sci. U. S. A. 2021, 118, e2021446118.

[14] C. Sun, B. Li, M. Zhao, S. Wang, Z. Lei, L. Lu, H. Zhang, L. Feng, C. Dou, D. Yin, H. Xu, Y. Cheng, F. Zhang, J. Am. Chem. Soc. 2019, 141, 19221.

[15] S. Liu, R. Chen, J. Zhang, Y. Li, M. He, X. Fan, H. Zhang, X. Lu, R. T. K. Kwok, H. Lin, J. W. Y. Lam, J. Qian, B. Z. Tang, ACS Nano 2020, 14, 14228.

[16] Y. Li, Z. Cai, S. Liu, H. Zhang, S. T. H. Wong, J. W. Y. Lam, R. T. K. Kwok, J. Qian, B. Z. Tang, Nat. Commun. 2020, 11, 1255.

[17] Y. Li, Y. Liu, Q. Li, X. Zeng, T. Tian, W. Zhou, Y. Cui, X. Wang, X. Cheng, Q. Ding, X. Wang, J. Wu, H. Deng, Y. Li, X. Meng, Z. Deng, X. Hong, Y. Xiao, Chem. Sci. 2020, 11, 2621.

[18] Y. Zhong, Z. Ma, F. Wang, X. Wang, Y. Yang, Y. Liu, X. Zhao, J. Li, H. Du, M. Zhang, Q. Cui, S. Zhu, Q. Sun, H. Wan, Y. Tian, Q. Liu, W. Wang, K. C. Garcia, H. Dai, Nat. Biotechnol. 2019, 37, 1322.

[19] M. Zhang, J. Yue, R. Cui, Z. Ma, H. Wan, F. Wang, S. Zhu, Y. Zhou, Y. Kuang, Y. Zhong, D. W. Pang, H. Dai, Proc. Natl. Acad. Sci. U. S. A. 2018, 115, 6590.

[20] Z. Ma, M. Zhang, J. Yue, C. Alcazar, Y. Zhong, T. C. Doyle, H. Dai, N. F. Huang, Adv. Funct. Mater. 2018, 28, 1803417.

[21] R. Tian, H. Ma, S. Zhu, J. Lau, R. Ma, Y. Liu, L. Lin, S. Chandra, S. Wang, X. Zhu, H. Deng, G. Niu, M. Zhang, A. L. Antaris, K. S. Hettie, B. Yang, Y. Liang, X. Chen, Adv. Mater. 2020, 32, 1907365.

[22] W. Wang, Z. Feng, B. Li, Y. Chang, X. Li, Y. Xu, R. Chen, X. Yu, H. Zhao, G. Lu, X. Kong, J. Qian, X. Liu, J. Mater. Chem. B, https://doi.org/10.1039/D0TB02728F.

[23] S. Wang, L. Liu, Y. Fan, A. M. El-Toni, M. S. Alhoshan, D. Li, F. Zhang, Nano. Lett. 2019, 19, 2418.

[24] Y. Zhong, Z. Ma, S. Zhu, J. Yue, M. Zhang, A. L. Antaris, J. Yuan, R. Cui, H. Wan, Y. Zhou, W. Wang, N. F. Huang, J. Luo, Z. Hu, H. Dai, Nat. Commun. 2017, 8, 737.

[25] Y. Li, S. Zeng, J. Hao, ACS Nano 2019, 13, 248.

[26] D. Chen, Y. Liu, Z. Zhang, Z. Liu, X. Fang, S. He, C. Wu, Nano. Lett. 2021, 21, 798.

[27] I. Gerhard, B. Monga, A. Waldbrenner, B. Runnebaum, J. Toxicol. Environ. Health A 1998, 54, 593.

[28] L. W. Jackson, P. P. Howards, J. Wactawski-Wende, E. F. Schisterman, Hum. Reprod. 2011, 26, 2887.

[29] F. D. S. Cordier, L. Mandereau, D. Hemon, Br. J. Ind. Med. 1991, 48, 375.

[30] J. Qian, B. Z. Tang, Chem 2017, 3, 56.

[31] D. Ding, K. Li, B. Liu, B. Z. Tang, Acc. Chem. Res. 2013, 46, 2441.

[32] W. Yu, B. Guo, H. Zhang, J. Zhou, X. Yu, L. Zhu, D. Xue, W. Liu, X. Sun, J. Qian, Science Bulletin 2019, 64, 410.

[33] W. Wu, Y. Yang, Y. Yang, Y. Yang, K. Zhang, L. Guo, H. Ge, X. Chen, J. Liu, H. Feng, Small 2019, 15, 1805549.

[34] Y. Wang, M. Chen, N. Alifu, S. Li, W. Qin, A. Qin, B. Z. Tang, J. Qian, ACS Nano 2017, 11, 10452.

[35] W. Liu, Y. Zhang, J. Qi, J. Qian, B. Z. Tang, Chem. Res. Chin. Univ. 2021, 37, 171.

[36] J. Qi, N. Alifu, A. Zebibula, P. Wei, J. W. Y. Lam, H. Q. Peng, R. T. K. Kwok, J. Qian, B. Z. Tang, Nano Today 2020, 34, 100893.

[37] Z. Zheng, D. Li, Z. Liu, H. Q. Peng, H. H. Y. Sung, R. T. K. Kwok, I. D. Williams, J. W. Y. Lam, J. Qian, B. Z. Tang, Adv. Mater. 2019, 31, 1904799.

[38] J. Qi, C. Sun, A. Zebibula, H. Zhang, R. T. K. Kwok, X. Zhao, W. Xi, J. W. Y. Lam, J. Qian, B. Z. Tang, Adv. Mater. 2018, 30, 1706856.

[39] W. Xu, D. Wang, B. Z. Tang, Angew. Chem. Int. Ed. 2020, 59, 2.

[40] C. Chen, H. Ou, R. Liu, D. Ding, Adv. Mater. 2020, 32, 1806331.

[41] H. Yan, Y. Xie, A. Wu, Z. Cai, L. Wang, C. Tian, X. Zhang, H. Fu, Adv. Mater. 2019, 31, 1901174.

[42] C. Chen, X. Ni, H. W. Tian, Q. Liu, D. S. Guo, D. Ding, Angew. Chem. Int. Ed. 2020, 59, 10008.

[43] X. Ni, X. Zhang, X. Duan, H. L. Zheng, X. S. Xue, D. Ding, Nano. Lett. 2018, 19, 318.

[44] N. Alifu, A. Zebibula, J. Qi, H. Zhang, C. Sun, X. Yu, D. Xue, J. W. Y. Lam, G. Li, J. Qian, B. Z. Tang, ACS Nano 2018, 12, 11282.

[45] Z. Feng, S. Bai, J. Qi, C. Sun, Y. Zhang, X. Yu, H. Ni, D. Wu, X. Fan, D. Xue, S. Liu, M. Chen, J. Gong, P. Wei, M. He, J. W. Y. Lam, X. Li, B. Z. Tang, L. Gao, J. Qian, (preprint) bioRxiv: 2020.05.26.113316, v2, submitted: Feb 2021.

[46] N. Alifu, A. Zebibula, H. Zhang, H. Ni, L. Zhu, W. Xi, Y. Wang, X. Zhang, C. Wu, J. Qian, Nano Res. 2020, 13, 2632.

[47] N. G. Horton, K. Wang, D. Kobat, C. G. Clark, F. W. Wise, C. B. Schaffer, C. Xu, Nat. Photonics 2013, 7, 205.

[48] L. Zhu, Y. Li, A. Xu, Hum. Reprod. 2012, 27, 2684.

[49] K. Van. Heertum, L. Barmat, Womens Health 2014, 10, 645.

[50] H. Leonhardt, B. Gull, K. Kishimoto, M. Kataoka, L. Nilsson, P. O. Janson, E. Stener-Victorin, M. Hellstrom, Acta Radiol. 2012, 53, 1195.

