## supporting information for "Aggregation-induced emission dots assisted non-invasive fluorescence hysterography in near-infrared IIb window"

***Scanning transmission electron microscopy (STEM).***

The AIE dots were negatively stained with uranyl acetate, then observed by Nova nano 450.

***Dynamic light scattering (DLS) measurement.***

The DLSs of AIE dots were measured by Zetasizer Nano-ZS.

***Synthesis of 4,4'-(5,6-dinitrobenzo[c][1,2,5]thiadiazole-4,7-diyl)bis(N,N-bis(4-(octyloxy)phenyl)aniline).***

4-(Octyloxy)-*N*-(4-(octyloxy)phenyl)-*N*-(4-(tributylstannyl)phenyl)aniline (1.58 g, 2 mmol), 4,7-dibromo-5,6-dinitrobenzo[*c*][1,2,5]thiadiazole (307 mg, 0.8 mmol), and $Pd(PPh_3)_4$ (47 mg, 0.04 mmol) were added into a 100 mL of two-necked round-bottom flask. The flask was then vacuumed and purged with dry nitrogen three times, and anhydrous THF (35 mL) was added. The mixture was heated to reflux and stirred for 24 h. After cooling down to room temperature, water was added, and the mixture was washed with dichloromethane three times. The organic phase was combined, dried with $MgSO_4$, and the solvent was evaporated under reduced pressure. The crude product was purified by column chromatography on silica gel using dichloromethane/hexane (v/v 1:2) as the eluent to afford 4,4'-(5,6-dinitrobenzo[*c*][1,2,5]thiadiazole-4,7-diyl)bis(*N*,*N*-bis(4-(octyloxy)phenyl)aniline) as a dark blue solid (73% yield). $^1$H NMR (400 MHz, $CDCl_3$): δ (ppm) 7.35 (d, 4H), 7.15 (d, 8H), 6.95 (d, 4H), 6.87 (d, 8H), 3.94 (t, 8H), 1.85–1.73 (m, 8H), 1.51–1.41 (m, 8H), 1.39–1.24 (m, 32H), 0.89 (t, 12H). $^{13}$C NMR (100 MHz, $CDCl_3$): δ (ppm) 156.49, 153.34, 150.63, 142.14, 139.25, 130.14, 127.91, 120.37, 117.73, 115.51, 68.28, 31.83, 29.38, 29.33, 29.26, 26.09, 22.68, 14.12.

***Synthesis of 4,7-bis(4-(bis(4-(octyloxy)phenyl)amino)phenyl)benzo[c][1,2,5]thiadiazole-5,6-diamine.***

To the mixture of 4,4'-(5,6-dinitrobenzo[*c*][1,2,5]thiadiazole-4,7-diyl)bis(*N*,*N*-bis(4-(octyloxy)phenyl)aniline) (613 mg, 0.5 mmol) and acetic acid (60 mL) in a 250 mL of two-necked round-bottom flask, iron powder (0.84 g, 15 mmol) was added. The mixture was heated to 80 °C, and stirred for 2 h. After cooling down to room temperature, water was added, and the mixture was washed with dichloromethane three times. The organic phase was combined, dried with $MgSO_4$, and the solvent was evaporated under reduced pressure. The crude product was purified by column chromatography on silica gel using dichloromethane/hexane (v/v 2:1) as the eluent to afford 4,7-bis(4-(bis(4-(octyloxy)phenyl)amino)phenyl)benzo[*c*][1,2,5]thiadiazole-5,6-diamine as a dark red solid (71% yield). $^1$H NMR (400 MHz, $CDCl_3$): δ (ppm) 7.36 (d, 4H), 7.15 (d, 8H), 7.06 (d, 4H), 6.85 (d, 8H), 4.09 (s, 4H), 3.94 (t, 8H), 1.82–1.74 (m, 8H), 1.50–1.41 (m, 8H), 1.35–1.26 (m, 32H), 0.90 (t, 12H). $^{13}$C NMR (100 MHz, $CDCl_3$): δ (ppm) 155.78, 151.39, 148.51, 140.38, 138.14, 130.74, 127.21, 119.72, 115.34, 113.64, 68.28, 31.83, 29.38, 29.36, 29.25, 26.09, 22.67, 14.11.

***Synthesis of 4,7-bis(4-(bis(4-(octyloxy)phenyl)amino)phenyl) benzo[1,2-c:4,5-c']bis[1,2,5]thiadiazole (OTPA-BBT).***

4,7-Bis(4-(bis(4-(octyloxy)phenyl)amino)phenyl)benzo[c][1,2,5]thiadiazole-5,6-diamine (350 mg, 0.3 mmol) was dissolved in dry pyridine (40 mL) in a 100 mL two-necked round-bottom flask. The flask was vacuumed and purged with dry nitrogen three times. Then N-sulfinylaniline (0.09 mL, 0.8 mmol) and trimethylsilyl chloride (0.15 mL, 1.2 mmol) were added into the solution. The mixture was heated to 80 °C, and stirred overnight. The solvent was evaporated under reduced pressure, and the residue was purified by column chromatography on silica gel using dichloromethane/hexane (v/v 1:2) as the eluent to result in 4,7-bis(4-(bis(4-(octyloxy)phenyl)amino)phenyl) benzo[1,2-c:4,5-c']bis[1,2,5]thiadiazole (OTPA-BBT) as a dark solid (68% yield). 1H NMR (400 MHz, CD2Cl2, 25 oC): δ (ppm) 8.09 (d, 4H), 7.18 (d, 8H), 7.05 (d, 4H), 6.89 (d, 8H), 3.96 (t, 8H), 1.84–1.73 (m, 8H), 1.52–1.42 (m, 8H), 1.42–1.22

(m, 32H), 0.89 (t, 12H). 13C NMR (100 MHz, CD2Cl2): δ (ppm) 158.16, 154.65, 151.20, 141.85, 134.63, 129.38, 128.62, 121.69, 120.14, 117.29, 70.26, 33.75, 31.28, 31.24, 31.17, 27.96, 24.58, 15.77. HRMS (MALDI-TOF, m/z) [M]+ calcd for C74H92O4N6S2, 1192.6621; found, 1192.6624.

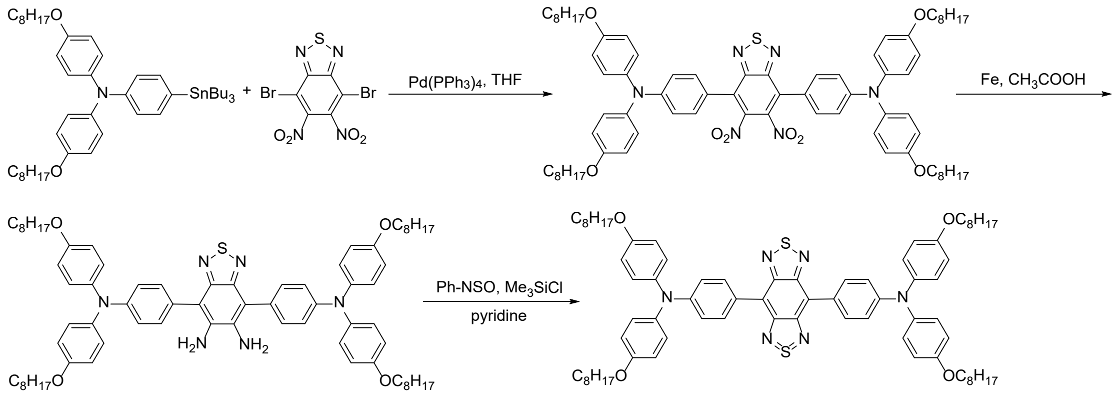


**Scheme S1.** Synthetic route to OTPA-BBT.

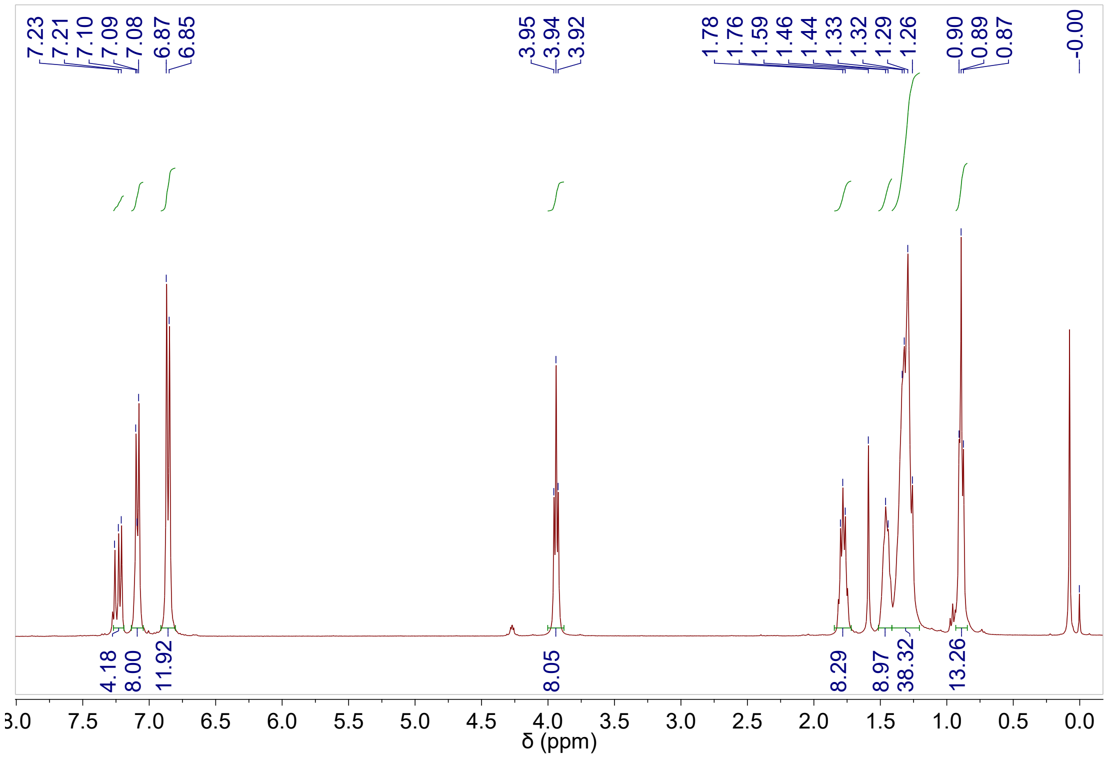


**Figure S1.** [1]H NMR spectrum of 4,4'-(5,6-dinitrobenzo[c][1,2,5]thiadiazole-4,7-diyl)bis(N,N-bis(4-(octyloxy)phenyl)aniline) in CDCl3 at 298 K.

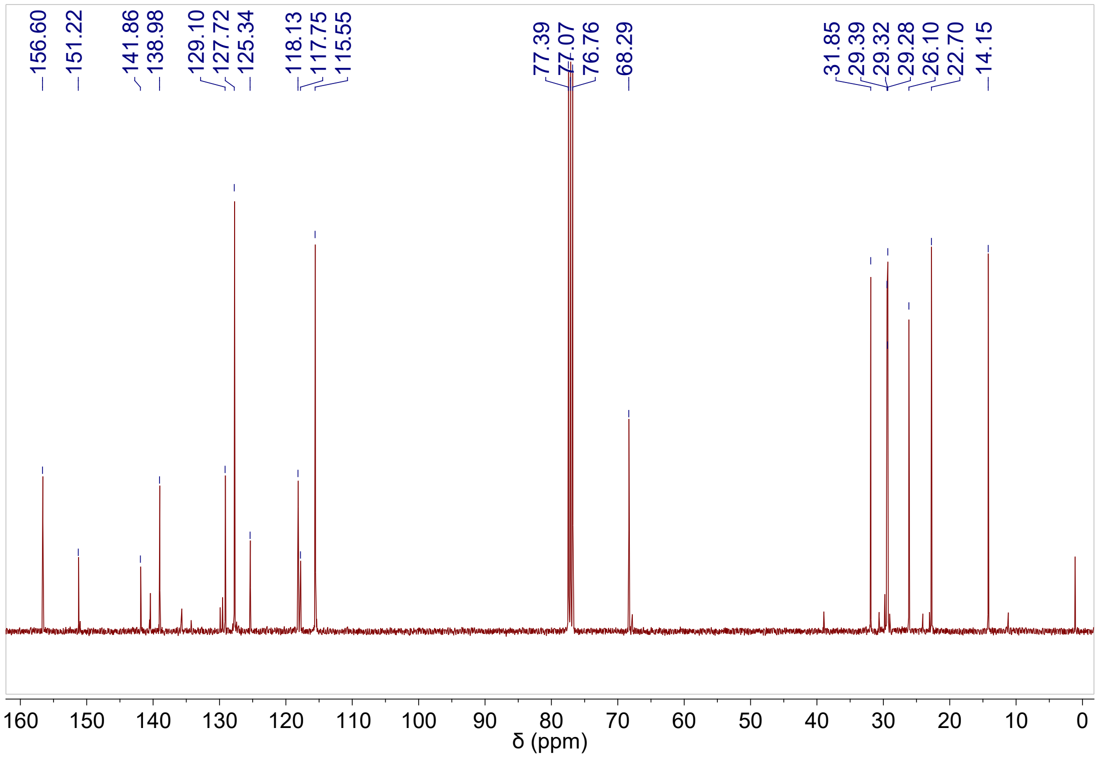


**Figure S2.** [13]C NMR spectrum of 4,4'-(5,6-dinitrobenzo[c][1,2,5]thiadiazole-4,7-diyl)bis(N,N-bis(4-(octyloxy)phenyl)aniline) in $CDCl_3$ at 298 K. C) HRMS of 4,4'-(5,6-dinitrobenzo[c][1,2,5]thiadiazole-4,7-diyl)bis(N,N-bis(4-(octyloxy)phenyl)aniline).

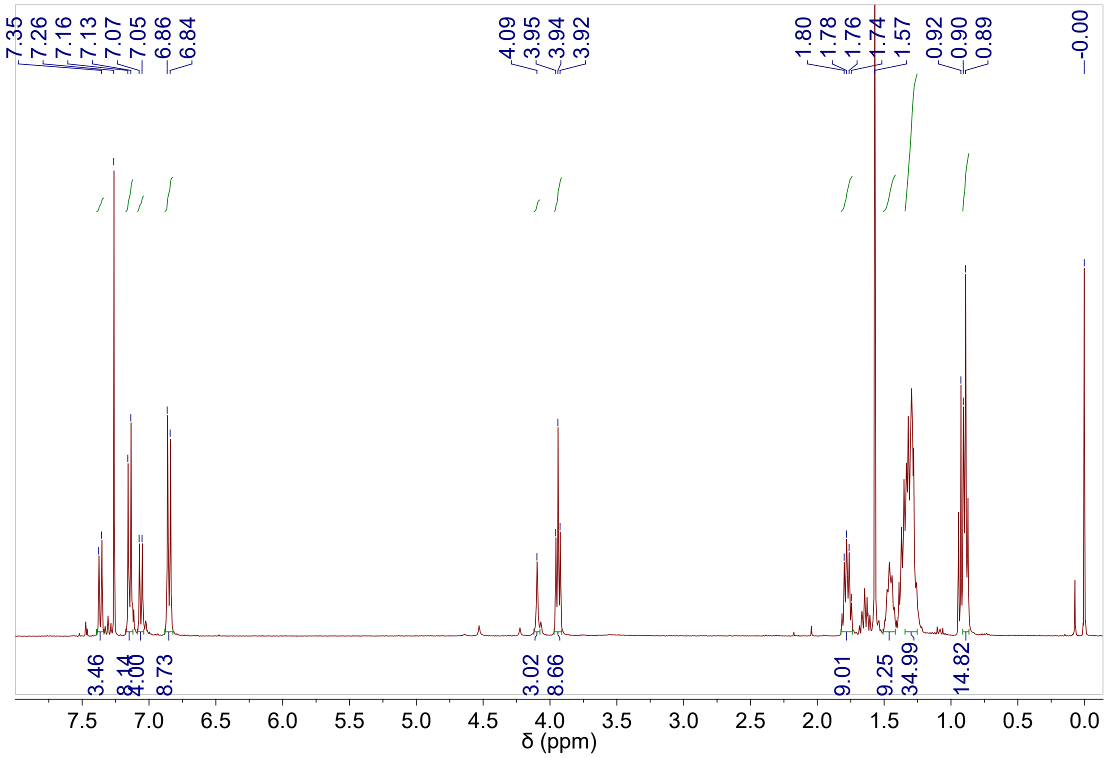

**Figure S3.** [1]H NMR spectrum of 4,7-bis(4-(bis(4-(octyloxy)phenyl)amino)phenyl)benzo[c][1,2,5]thiadiazole-5,6-diamine in $CDCl_3$ at 298 K.

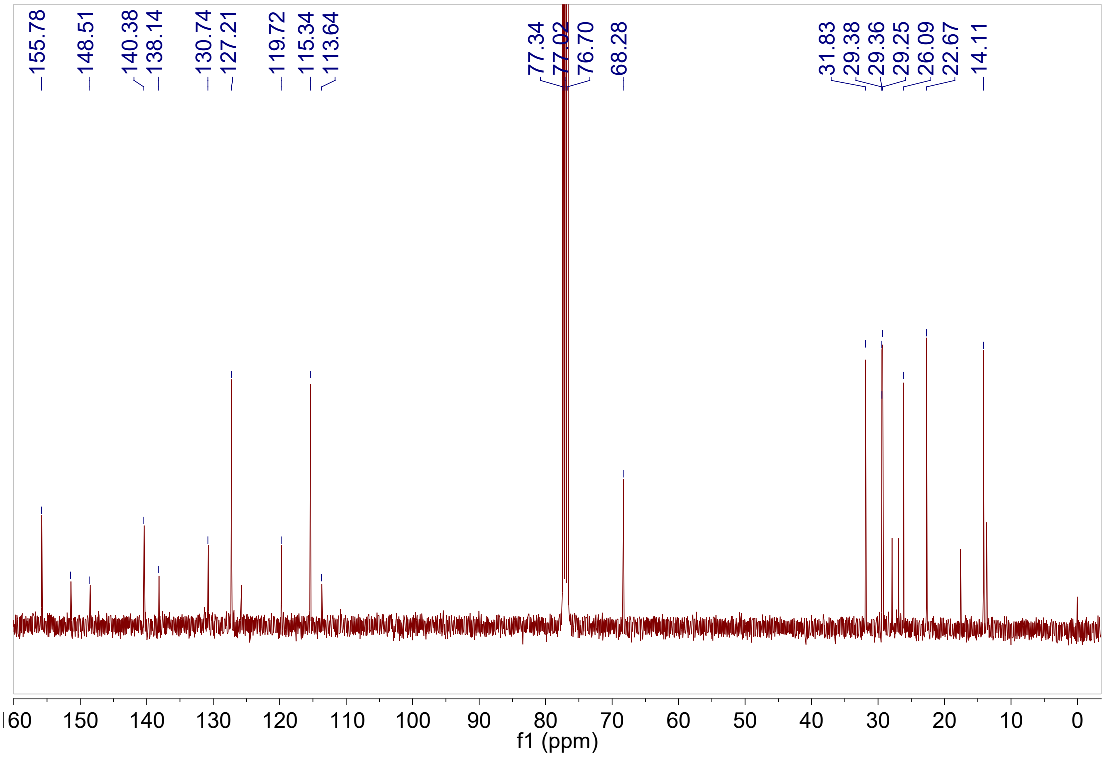


**Figure S4.** [13]C NMR spectrum of 4,7-bis(4-(bis(4-(octyloxy)phenyl)amino)phenyl)benzo[c][1,2,5]thiadiazole-5,6-diamine in $CDCl_3$ at 298 K.

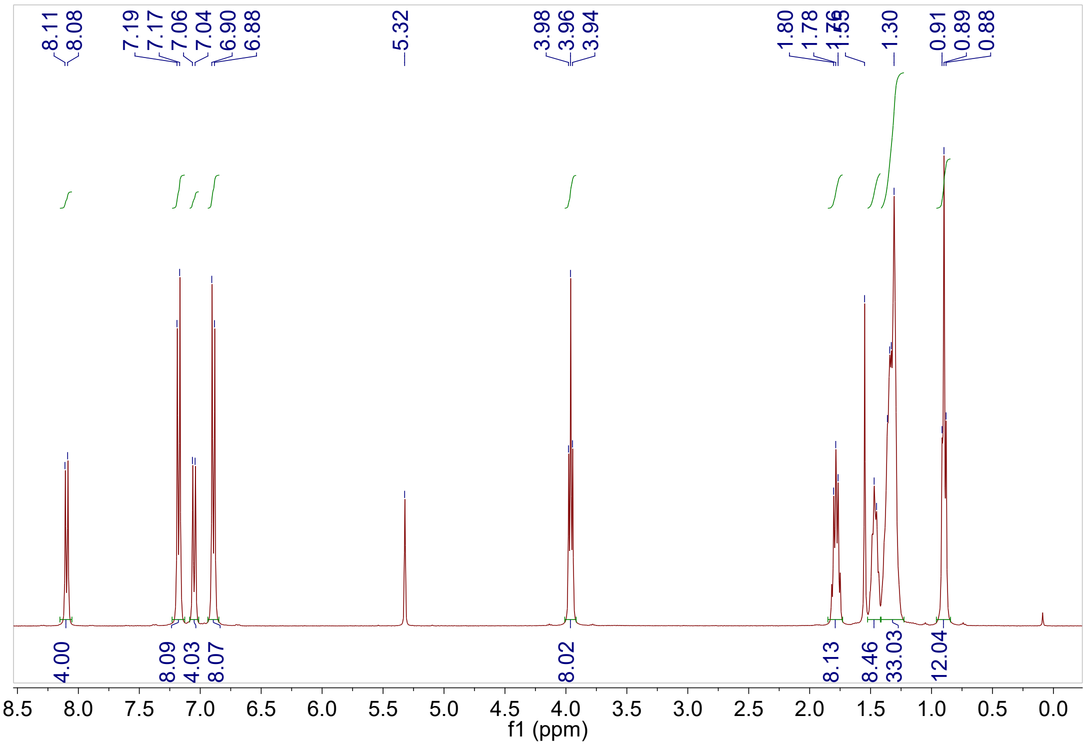


**Figure S5.** $^{1}$H NMR spectrum of OTPA-BBT in $CD_2Cl_2$ at 298 K.

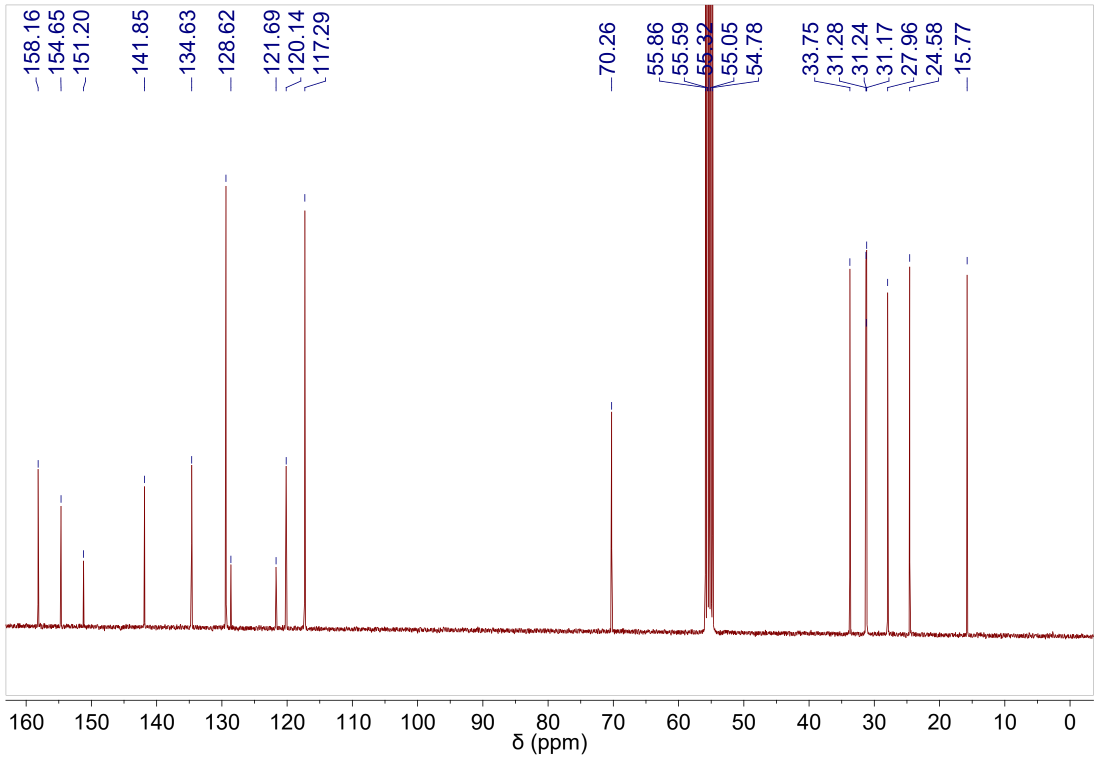


**Figure S6.** $^{13}$C NMR spectrum of OTPA-BBT in $CD_2Cl_2$ at 298 K.

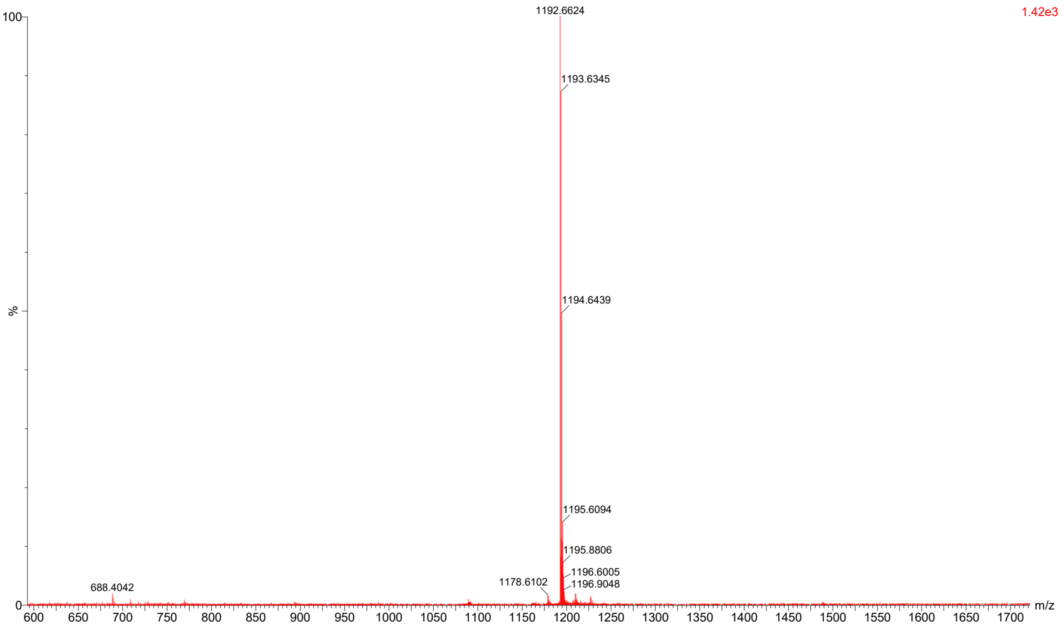


**Figure S7.** HRMS of OTPA-BBT.

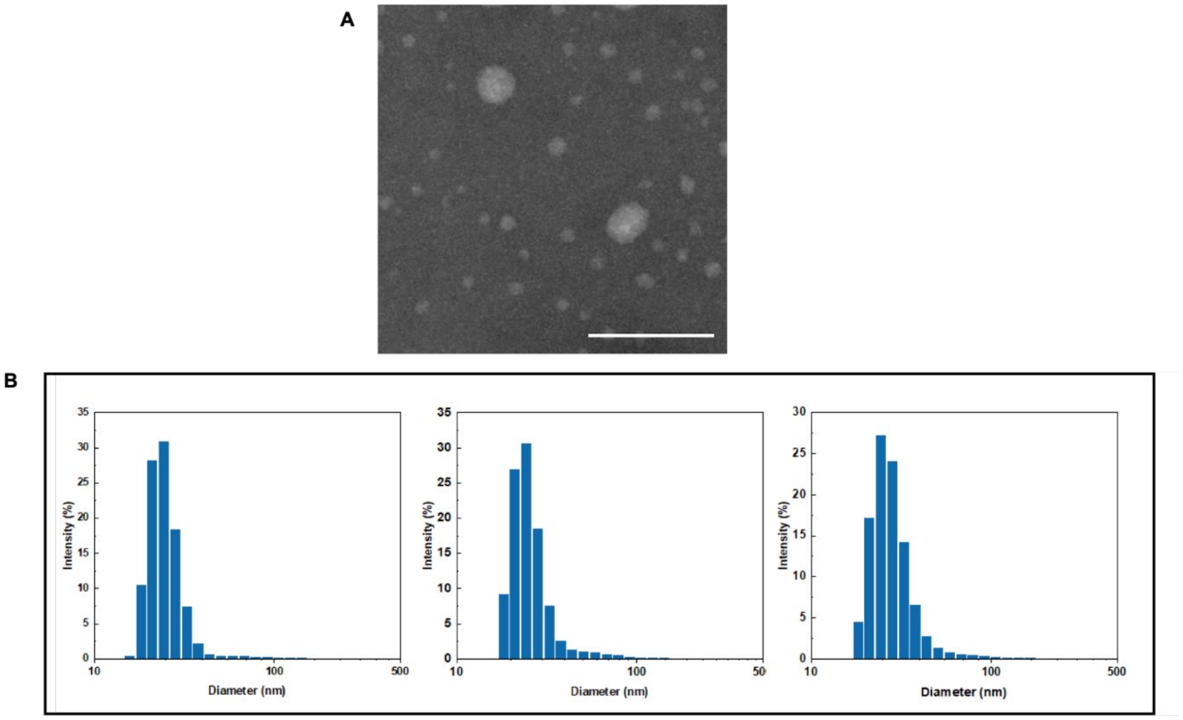


**Figure S8.** A) STEM image and B) DLS results (28.3 ± 1.6 nm) of OTPA-BBT dots. Scale bar, 100 nm.

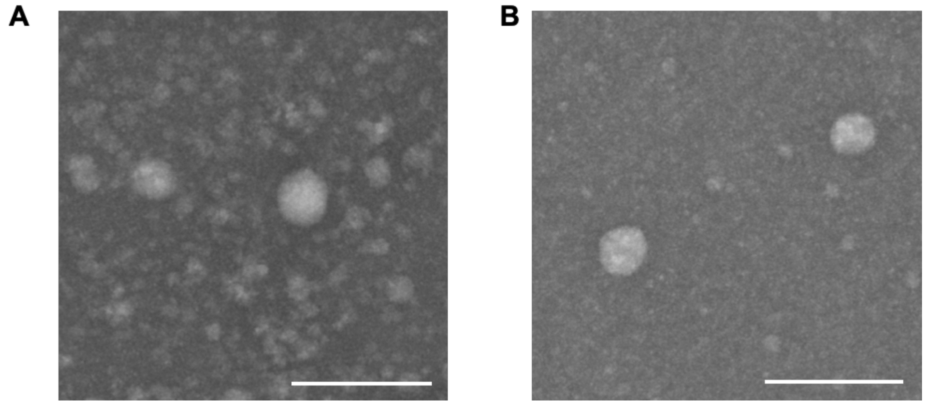


**Figure S9.** STEM images of OTPA-BBT dots dissolved in A) acidic solution (pH = 2) and B) alkaline solution (pH = 12). Scale bar, 100 nm.

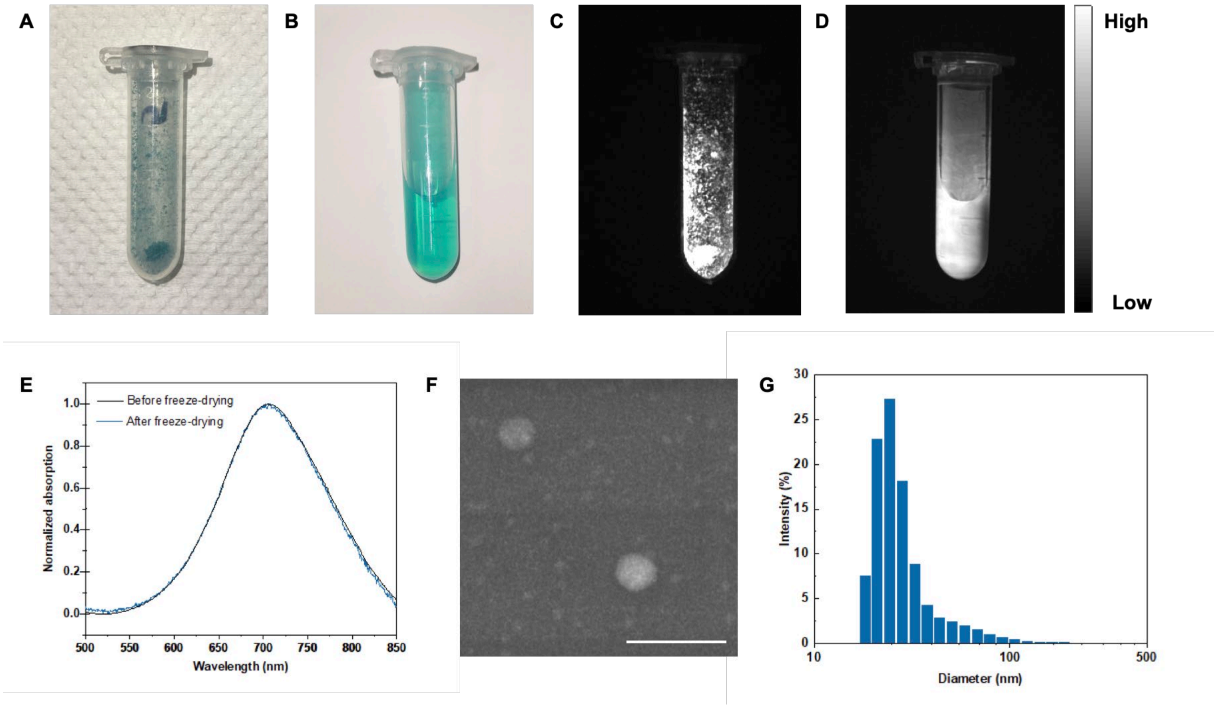


**Figure S10.** Photophysical properties of OTPA-BBT dots after freeze-drying and rehydration. A) Lyophilized OTPA-BBT dots. B) Solution of lyophilized OTPA-BBT dots. C) NIR-IIb fluorescence image of freeze-dried OTPA-BBT dots. D) NIR-IIb fluorescence image of rehydrated OTPA-BBT dots. G) The normalized absorption spectra of OTPA-BBT dots before freeze-drying and rehydrated OTPA-BBT dots. E) Lyophilized OTPA-BBT dots observed by scanning transmission electron microscope. Scale bar, 100 nm. G) The DLS result of the rehydrated OTPA-BBT dots (mean size = 29.6 nm).

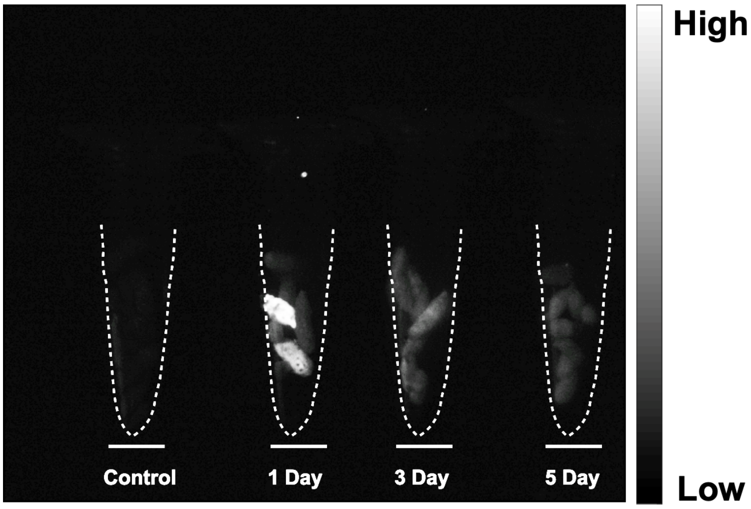


**Figure S11.** NIR-II fluorescence fecal image (beyond 1300 nm) under 793 nm excitation after treatment. The power density of the laser irradiation was ~18 mW cm$^{-2}$ and the exposure time was 100 ms.

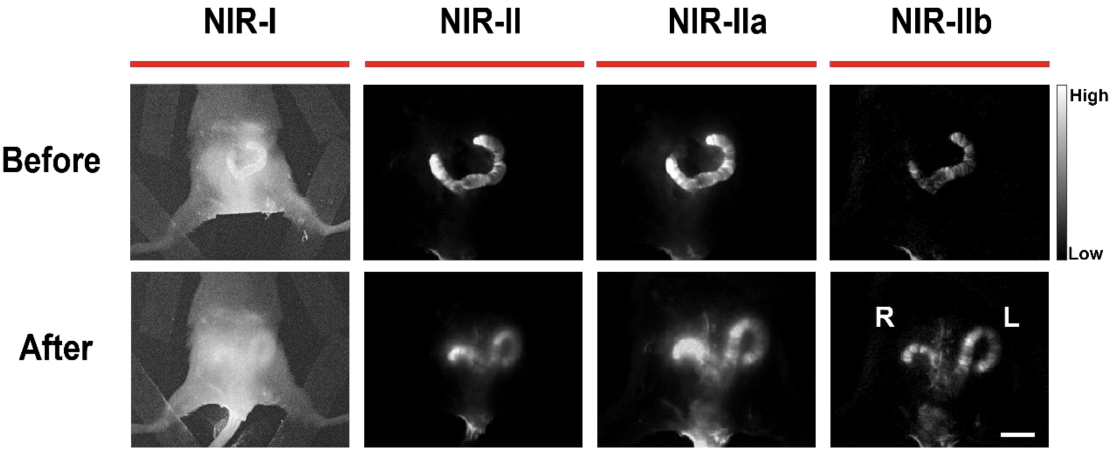


**Figure S12.** The comparison of the NIR-I (800-900 nm), NIR-II (1000-1700 nm), NIR-IIa (1300-1400 nm) and NIR-IIb (1500-1700 nm) fluorescence uterine imaging in the mouse before and after closing the abdominal cavity. Scale bar, 10 mm. L, left; R, right. The power densities of the laser irradiation were ~20, ~70, ~100, and ~100 mW cm$^{-2}$, respectively. The integrated times were 7.7, 1, 50, and 100 ms, respectively.

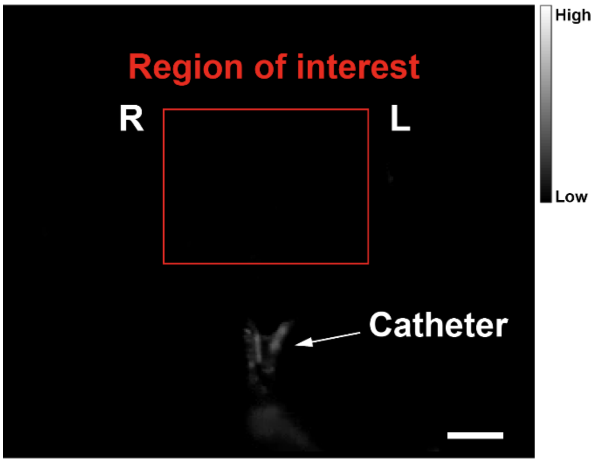


**Figure S13**. The NIR-IIb fluorescence image after flushing the abdominal cavity and the fluorescent dots flowing away along the cervix and vagina. Scale bar, 10 mm. L, left; R, right. The power density of the laser irradiation was ~80 mW cm$^{-2}$ and the integrated time was 150 ms.

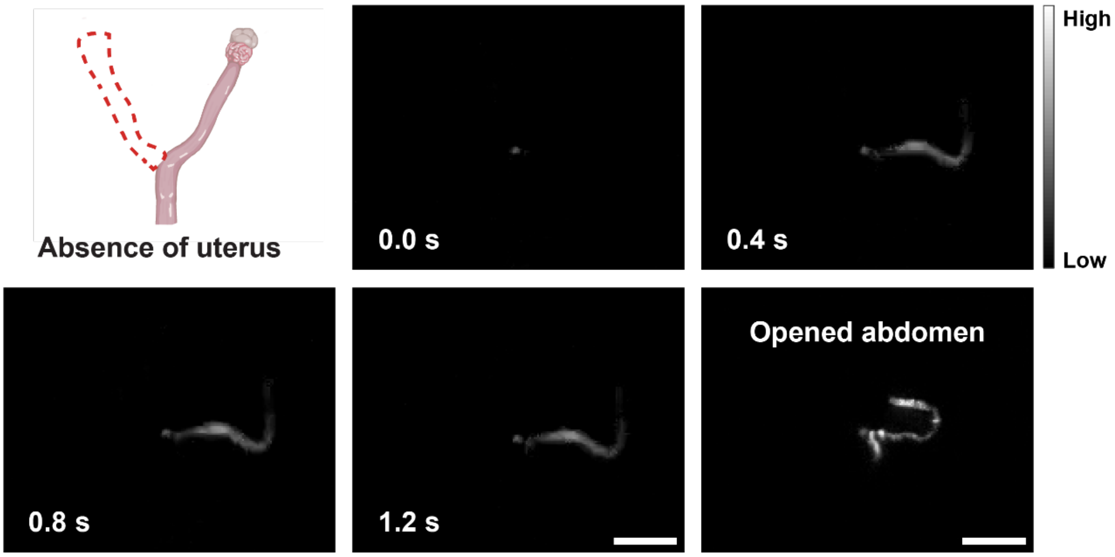


**Figure S14.** The observation of the absence of the uterus with time. The illustration in the top left hand corner briefly explained the disease model. The NIR-IIb image in the bottom right hand corner was taken after opening the abdominal cavity. Scale bar, 10 mm. The power density of the laser irradiation was ~80 mW cm$^{-2}$ and the integrated time was 150 ms.

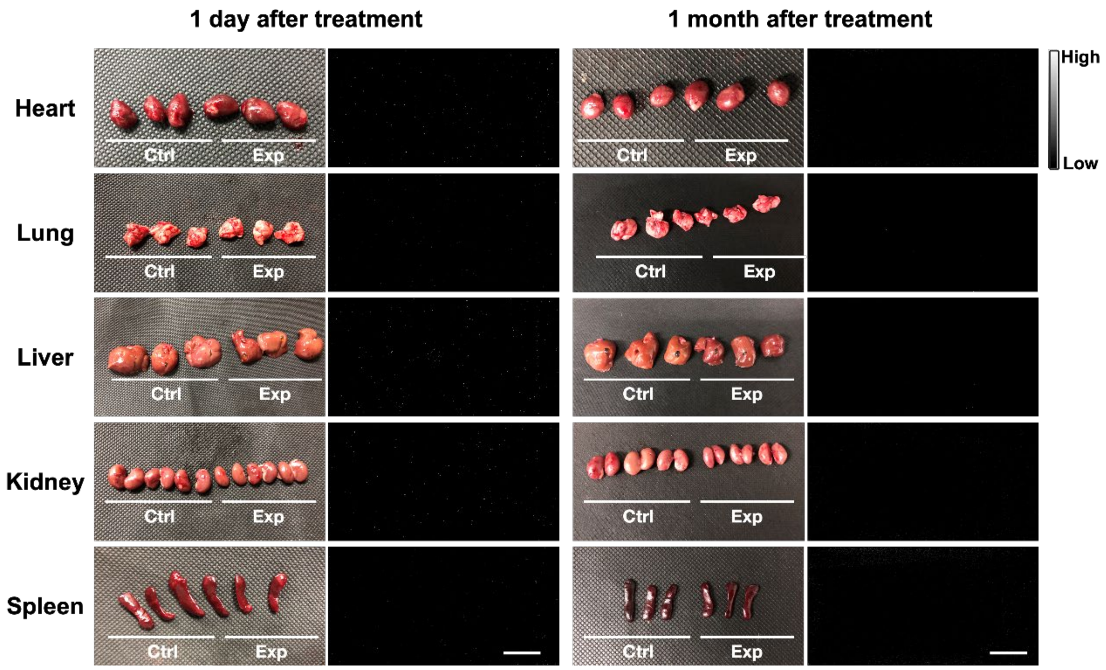


**Figure S15.** The observation of main organs after treatment. At 1 day and 1 month after intrauterine perfusion, no fluorescence signal was observed in NIR-II fluorescence (beyond 1300 nm) organ imaging. Scale bar, 30 mm. The power density of the laser irradiation was ~80 mW cm$^{-2}$ and the exposure time was 150 ms.

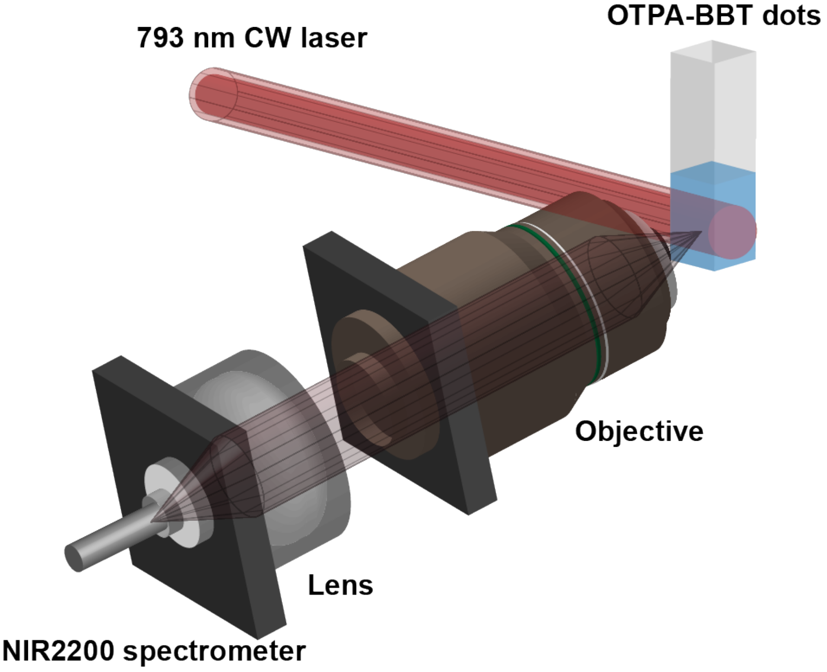


**Figure S16.** Schematic illustration of the NIR-II fluorescence spectra measurement system.

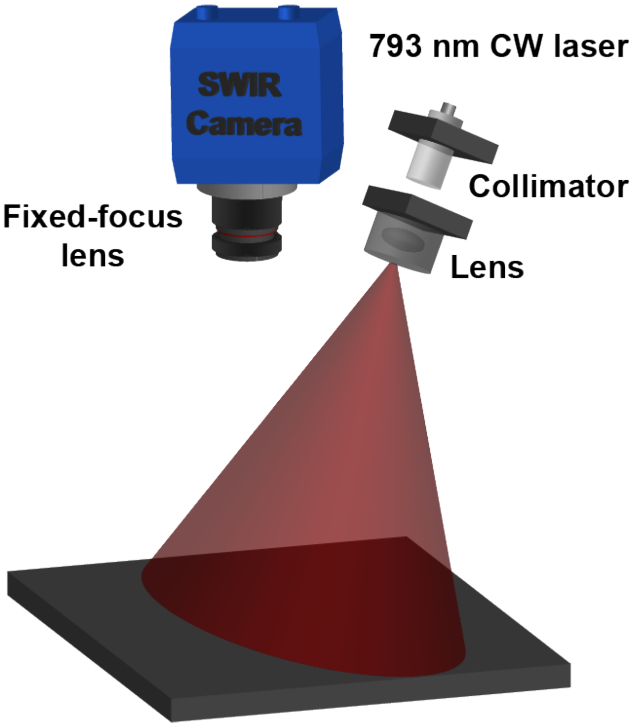


**Figure S17.** Schematic illustration of the NIR-II fluorescence macroscopic imaging system.

**Movie S1.** Dynamic fluorescence hysterography recording the uterus expelling fluid through peristalsis.

**Movie S2**. OTPA-BBT dots filling the cavity and then spilling from the ruptured uterus.

**Movie S3.** Real-time fluorescence hysterography of the uterus after surgical suture.
